# *Wolbachia pipientis* associated to tephritid fruit fly pests: from basic research to applications

**DOI:** 10.1101/358333

**Authors:** Mariana Mateos, Humberto Martinez, Silvia B. Lanzavecchia, Claudia Conte, Karina Guillén, Brenda M. Morán-Aceves, Jorge Toledo, Pablo Liedo, Elias D. Asimakis, Vangelis Doudoumis, Georgios A. Kyritsis, Nikos T. Papadopoulos, Antonios A. Avgoustinos, Diego F. Segura, George Tsiamis, Kostas Bourtzis

## Abstract

Members of the true fruit flies (family Tephritidae) are among the most serious agricultural pests worldwide, whose control and management demands large and costly international efforts. The need for cost-effective and environmentally-friendly integrated pest management (IPM) has led to the development and implementation of autocidal control strategies. Autocidal approaches include the widely used sterile insect technique (SIT) and the incompatible insect technique (IIT). IIT relies on maternally transmitted bacteria (namely *Wolbachia*), to cause a conditional sterility in crosses between released mass-reared *Wolbachia*-infected males and wild females, which are either uninfected or infected with a different *Wolbachia* strain (i.e., cytoplasmic incompatibility; CI). Herein, we review the current state of knowledge on *Wolbachia*-tephritid interactions including infection prevalence in wild populations, phenotypic consequences, and their impact on life history traits. Numerous pest tephritid species are reported to harbor *Wolbachia* infections, with a subset exhibiting high prevalence. The phenotypic effects of *Wolbachia* have been assessed in very few tephritid species, due in part to the difficulty of manipulating *Wolbachia* infection (removal or transinfection). Based on recent methodological advances (high-throughput DNA sequencing) and a breakthrough concerning the mechanistic basis of CI, we suggest research avenues that could accelerate generation of necessary knowledge for the potential use of *Wolbachia*-based IIT in area-wide integrated pest management (AW-IPM) strategies for the population control of tephritid pests.

## 1 Background

### 1.1 The economic importance of tephritids as pests

Flies in the family Tephritidae (Diptera) include some of the world’s most important agricultural pests. The family is comprised of ˜4900 described species within 481 genera, of which seven (*Anastrepha*, *Bactrocera*, *Ceratitis*, *Dacus*, *Rhagoletis*, *Toxotrypana* and *Zeugodacus*) contain ˜70 major pest species [1–5]. In addition to causing billions of dollars in direct losses to a wide variety of fruit, vegetable and flower crops, pest tephritids limit the development of agriculture in many countries due to strict quarantines implemented in fruit-buying countries, and to the costs associated with efforts aimed at prevention, containment, suppression, and eradication.

To prevent or minimize the harmful effects of tephritid pests, growers of affected crops must comply with health and safety standards required by the market, applying an area-wide management approach involving chemical, biological, cultural, and autocidal control practices [6, 7]. Autocidal refers to methods that use the insect to control itself, by releasing insects that are sterile or induce sterility upon mating with wild insects in the next or subsequent generations [8–10]. Autocidal strategies include the sterile insect technique (SIT); one of the most widespread control methods used against fruit flies. SIT relies on the mass-rearing production, sterilization and recurrent release of insects (preferentially males) of the targeted species. Sterilization is attained by radiation, in a way that does not impair male mating and insemination capabilities. Wild females that mate with sterilized males lay unfertilized eggs. At the appropriate sterile:wild (S:W) ratio, the reproductive potential of the target population can be reduced [11–13]. Historically, at least 28 countries have used the SIT at a large-scale for the suppression or eradication of pest insects [14–16]. SIT has been applied successfully for several non-tephritid insect pests, for example: the New World screw worm (*Cochliomyia hominivorax* Coquerel); several species of tsetse fly (*Glossina* spp.); and the codling moth (*Cydia pomonella* L.) [reviewed in 17, 18].

Successful SIT programs as part of Area-wide Integrated Pest Management (IPM) strategies have also been implemented for several tephritids: *Ceratitis capitata* Wiedemann; *Anastrepha ludens* Loew; *Anastrepha obliqua* Macquart; *Anastrepha fraterculus* Wiedemann; *Zeugodacus cucurbitae* Coquillett; *Bactrocera dorsalis* Hendel; and *Bactrocera tryoni* Froggatt [6, 12, 13, 15]. SIT is currently being developed for two additional tephritid species: *Dacus ciliatus* Loew and *Bactrocera tau* Walker [19, 20]. The advantages of the SIT over other pest control approaches (e.g. use of pesticides) are that it is the most environmentally friendly and resistance is unlikely to evolve [21, 22].

Another autocidal strategy where mating between mass-reared and wild insects can be used to suppress pest populations is the incompatible insect technique (IIT). IIT also relies on the principle of reducing female fertility, but utilizes endosymbiotic bacteria instead of radiation, to induce a context-dependent sterility in wild females. It is based on the ability of certain maternally inherited bacteria (namely from the genus *Wolbachia*) to induce a form of reproductive incompatibility known as cytoplasmic incompatibility (CI; explained in the section below). Herein we review the current knowledge on taxonomic diversity of *Wolbachia*-tephritid associations and their phenotypic consequences, and identify gaps in knowledge and approaches in the context of potential application of IIT in AW-IPM programs to control tephritid pests. We also discuss scenarios where these two autocidal strategies, SIT and IIT, could be potentially combined for the population suppression of tephritid pests.

### 1.2 The influence of Wolbachia on host ecology

Insects and other arthropods are common hosts of maternally inherited bacteria [reviewed in 23]. These heritable endosymbionts can have a strong influence on host ecology. Such vertically transmitted bacteria are typically vastly (or fully) dependent on the host for survival and transmission. Certain associations are obligate for both partners, and generally involve a nutritional benefit to the host. Other heritable bacteria are facultative, with such associations ranging from mutualistic to parasitic from the host’s perspective. Among these, *Wolbachia* is the most common and widespread facultative symbiont of insects and arthropods [24–27].

*Wolbachia* is a diverse and old genus [possibly older than 200 million years; 28] of intracellular gram-negative Alphaproteobacteria (within the order Rickettsiales) associated with arthropods and filarial nematodes. *Wolbachia* cells resemble small spheres 0.2–1.5 μm, occur in all tissue types, but tend to be more prevalent in ovaries and testicles of infected hosts, and are closely associated with the female germline [reviewed by 29, see also 30]. *Wolbachia* is estimated to infect 40–66% of insect species [24–27]. Within a species or population, the infection prevalence of *Wolbachia* can be quite variable over space [e.g. 31] and time [e.g. 32, 33].

The most commonly documented effects of *Wolbachia* on arthropod hosts fall under the category of reproductive parasitism, which involves manipulation of host reproduction to enhance symbiont transmission and persistence, in general by increasing the relative frequency of *Wolbachia*-infected vs. uninfected females. Females are typically the sex that can transmit *Wolbachia* and other heritable bacteria, although rare exceptions exist [34–36]. *Wolbachia* employs all four types of reproductive manipulation [reviewed by 37, 38, 39]. **Feminization** results in genetic males that develop and function as females,and occurs in the orders Hemiptera, Lepidoptera, and Isopoda. *Wolbachia*-induced **parthenogenesis** occurs in haplo-diploid hosts (e.g. Acari, Hymenoptera and Thysanoptera), where unfertilized eggs, which would otherwise develop into males, develop into females. **Male killing** eliminates infected males to the advantage of surviving infected female siblings, and occurs in Coleoptera, Diptera, Lepidoptera, and Pseudoscorpiones. **Cytoplasmic incompatibility (CI)** [40] prevents infected males from producing viable offspring upon mating with females lacking *Wolbachia* (or a compatible strain of *Wolbachia*; see below). CI is the most commonly reported *Wolbachia*-induced reproductive phenotype, and is found in Acari, Coleoptera, Diptera, Hemiptera, Hymenoptera, Isopoda, Lepidoptera, and Orthoptera. In **unidirectional CI**, eggs from uninfected females that are fertilized by sperm from *Wolbachia*-infected males fail to develop (Fig. 1a). **Bi-directional CI** results from crosses involving two different (incompatible) *Wolbachia* strains (Fig. 1b). In contrast, crosses between females and males infected with the same or compatible *Wolbachia* strains are viable. Similarly, *Wolbachia*-infected females are compatible with uninfected males [39]. Consequently, above a certain threshold of *Wolbachia* infection frequency in a host population, infected females are at a reproductive advantage over uninfected females, resulting in a positive feedback loop that leads to the increase of *Wolbachia* frequencies [41].

**Figure 1.**
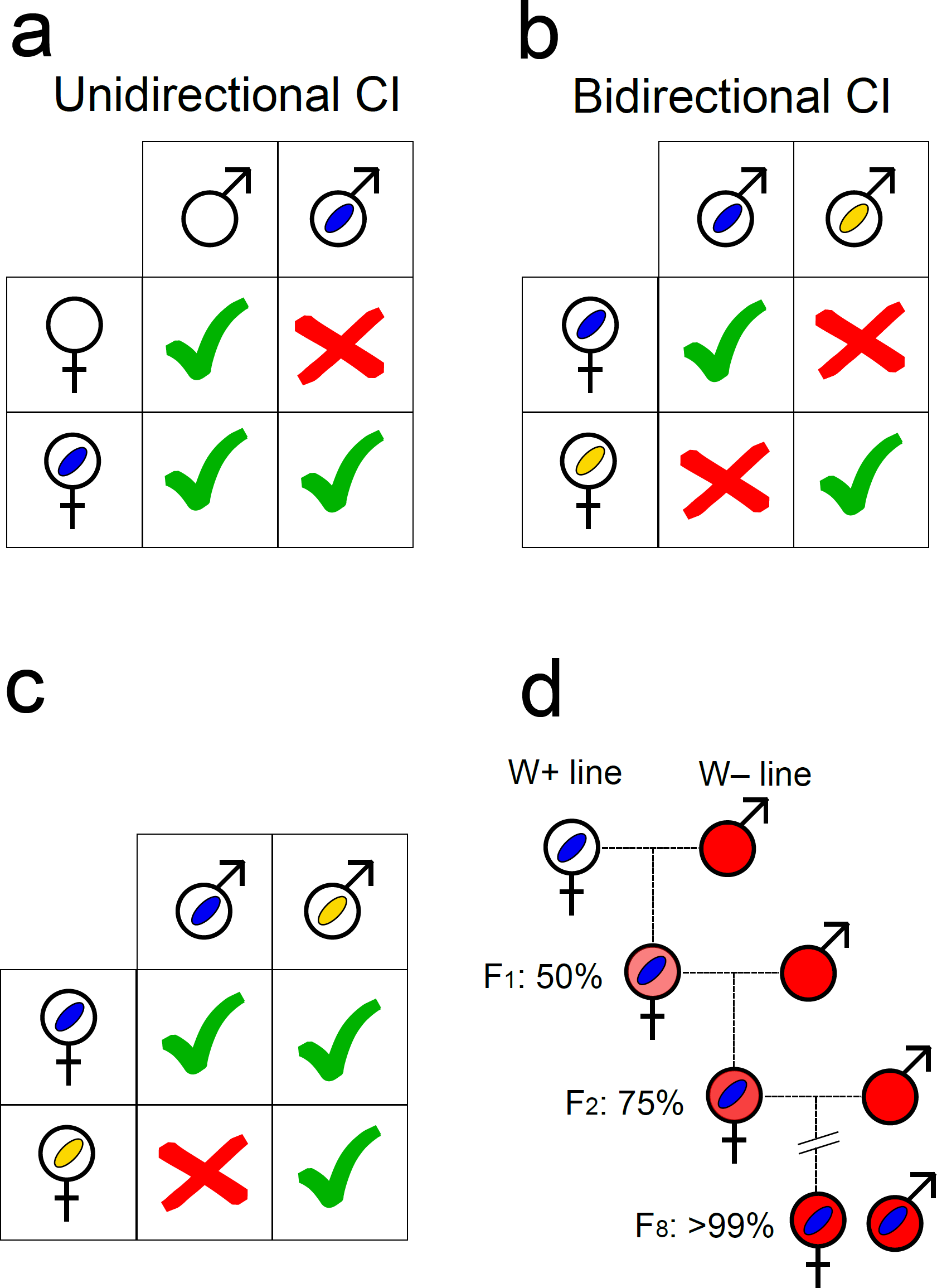
**a,b**. Qualitative illustration of uni- and bidirectional cytoplasmic incompatibility (CI) on the basis of the *Wolbachia* infection status of the parent generation. Empty male and female symbols signify absence of *Wolbachia*. Blue and yellow ovals represent distinct *Wolbachia* strains. Green tick marks = Successful offspring production. Red crosses = no offspring production. **c.** A special case of unidirectional incompatibility in which one *Wolbachia* strain (i.e., the blue one; equivalent to *w*Ri; see text) can rescue another strain (i.e., the yellow one; equivalent to *w*Mel), but not vice versa. **d.** Backcrossing procedure. *Wolbachia* infection is indicated by blue oval. Host nuclear backgrounds are indicated by colors: white represents the initial nuclear background of *Wolbachia*-infected host; red (darkest) indicates the host background of the *Wolbachia-*free line contributing males every generation. Different shades of red represent the increasing replacement of “white” nuclear background over backcrossing generations (F_1_to F_8_) by “red” nuclear background.

CI was discovered almost half a century ago [40], but its mechanism has not been fully elucidated. A useful conceptual model to understand the observed patterns of CI is “mod/resc” [42, 43]. It postulates that *Wolbachia* has two functions: *mod* (for modification), which acts as a toxin or imprint of the male germline; and *resc* (for rescue), which acts as an antidote. The *mod* function acts on the nucleus in the male germline, before *Wolbachia* are shed from maturing sperm [44]. When a sperm nucleus affected by *mod* enters the egg of an uninfected female, this nucleus encounters problems such as delays in DNA replication and cell-cycle progression, leading to embryo death. In contrast, if the appropriate *resc* (“the antidote”) function is active in the egg, the defect caused by *mod* in the sperm is rescued, and the embryo proceeds through normal development. Based on the mod/resc model, four different CI-*Wolbachia* types (strains) are possible: (1) the mod^+^/resc^+^ type, capable of inducing CI and rescuing its own modification; (2) the mod^−^/resc^−^ type, incapable of inducing CI or rescuing the CI effect of other mod^+^ strains; (3) the mod^+^/resc^−^ (“suicidal”) type, capable of inducing CI but not rescuing its own modification; and (4) the mod^−^/resc^+^ type, incapable of inducing CI, but able to rescue the CI defect caused by a mod^+^ strain [45, 46].

Three recent breakthrough studies identified a *Wolbachia*-encoded operon (of viral origin) that appears to be required to induce CI [47–49]. LePage et al. [47] used a combination of genomic, bioinformatic and molecular approaches to identify two contiguous genes (*cifA* and *cifB*) encoded in the genome of *Wolbachia w*Mel strain (from *D. melanogaster*). LePage et al. [47] found that homologous genes are present in CI-inducing strains, but are absent, or highly diverged, in non-CI-inducing strains. In addition, LePage et al. [47] found that strain *w*Ri (from *D. simulans*), which is compatible with strain *w*Mel, but not vice versa (see Fig. 1c) [50, 51], shares two sets of similar CI operons with *w*Mel, but has an additional divergent CI operon, which could explain why *w*Mel is unable to rescue the *w*Ri-induced defects. LePage et al. [47] then expressed *cifA* and *cifB* as transgenes in the germ line of *D. melanogaster* males. When these males were mated to *Wolbachia*-free females, the offspring had reduced hatch rates and embryonic cell-division defects that resembled those observed in CI. In contrast, when these males were mated to *Wolbachia*-infected females, hatching rate was restored. In a parallel study, Beckmann et al. [48] identified what appear to be two homologous CI operons in the strain *w*Pip (from *Culex pipiens* L.) named *cidA-cidB* and *cinA-cinB*. They also expressed them in *D. melanogaster* male germline, and found that when these males were mated to *Wolbachia*-free females, the offspring exhibited the characteristic embryonic abnormalities of CI. Based on the observation that the expression of *cidB* or *cinB* in yeast blocks growth, whereas expression of *cidA* or *cinA*, respectively rescue the defect, Beckmann et al. [48] proposed that *cidA* and *cinA* function as the antidote to *cidB* or *cinB*, respectively. A similar rescue phenomenon was not recapitulated in the fly, though. More recently, Shropshire et al. [49] demonstrated that *cifA* in *D. melanogaster* acts as the rescue gene, and thus propose a “Two-by-One” model whereby *cifA* and *cifB* induce CI and *cifA* rescues CI.

In addition to its reproductive phenotypes on arthropods, *Wolbachia* engages in obligate mutualistic interactions with filarial nematodes [39] and with members of five insect orders [reviewed in 52]. For example, it serves as a nutritional mutualist of bedbugs [53], and as an enabler of oogenesis in a parasitic wasp [54]. As a facultative symbiont, *Wolbachia* can provide direct fitness benefits to its insect hosts by influencing development, nutrition, iron metabolism, lifespan, and fecundity [55–60], and most notably, by conferring resistance or tolerance to pathogens (particularly single-stranded RNA viruses) and parasites [61–73]. The interference of *Wolbachia* with the replication and transmission of certain viruses forms the basis of several regional mosquito-borne disease control programs [74; e.g. http://www.eliminatedengue.com, 75, 76].

Certain host-*Wolbachia* combinations incur fitness costs to the host, beyond reproductive parasitism, including reduced longevity, sperm competitive ability, and fecundity, as well as higher susceptibility to natural enemies [77–84]. Similarly, certain host-*Wolbachia* combinations may potentially enhance pathogen-vectoring capacities [85–88].

## 2 Methods to study Wolbachia

### 2.1 Methods to assess Wolbachia infection status

For purposes of this review, we consider a host species or population as “infected” with *Wolbachia*, even if the infection is transient or found at low titer. *Wolbachia,* and most cytoplasmically transmitted endosymbionts, are fastidious to culture outside host cells, such that their study typically relies on culture-independent methods. A recommended flow-chart of steps is depicted in Fig. 2. The most utilized approach to date for identifying hosts infected with *Wolbachia* is through PCR screening of *Wolbachia* genes in DNA extracts of hosts. Different PCR primers have been used to perform such surveys, traditionally targeting a portion of the 16S ribosomal (r)RNA gene or of a ubiquitous protein-coding gene (e.g. *wsp* or *ftsZ*). Simoes et al. [89] evaluated the relative sensitivity and specificity of different primer pairs aimed at *Wolbachia* detection and identification, revealing that no single PCR protocol is capable of specific detection of all known *Wolbachia* strains. A related method known as “loop mediated isothermal amplification” (LAMP; not shown in Fig. 2), which requires less infrastructure than PCR, has been successfully employed for *Wolbachia* detection in several insects [90].

**Figure 2.**
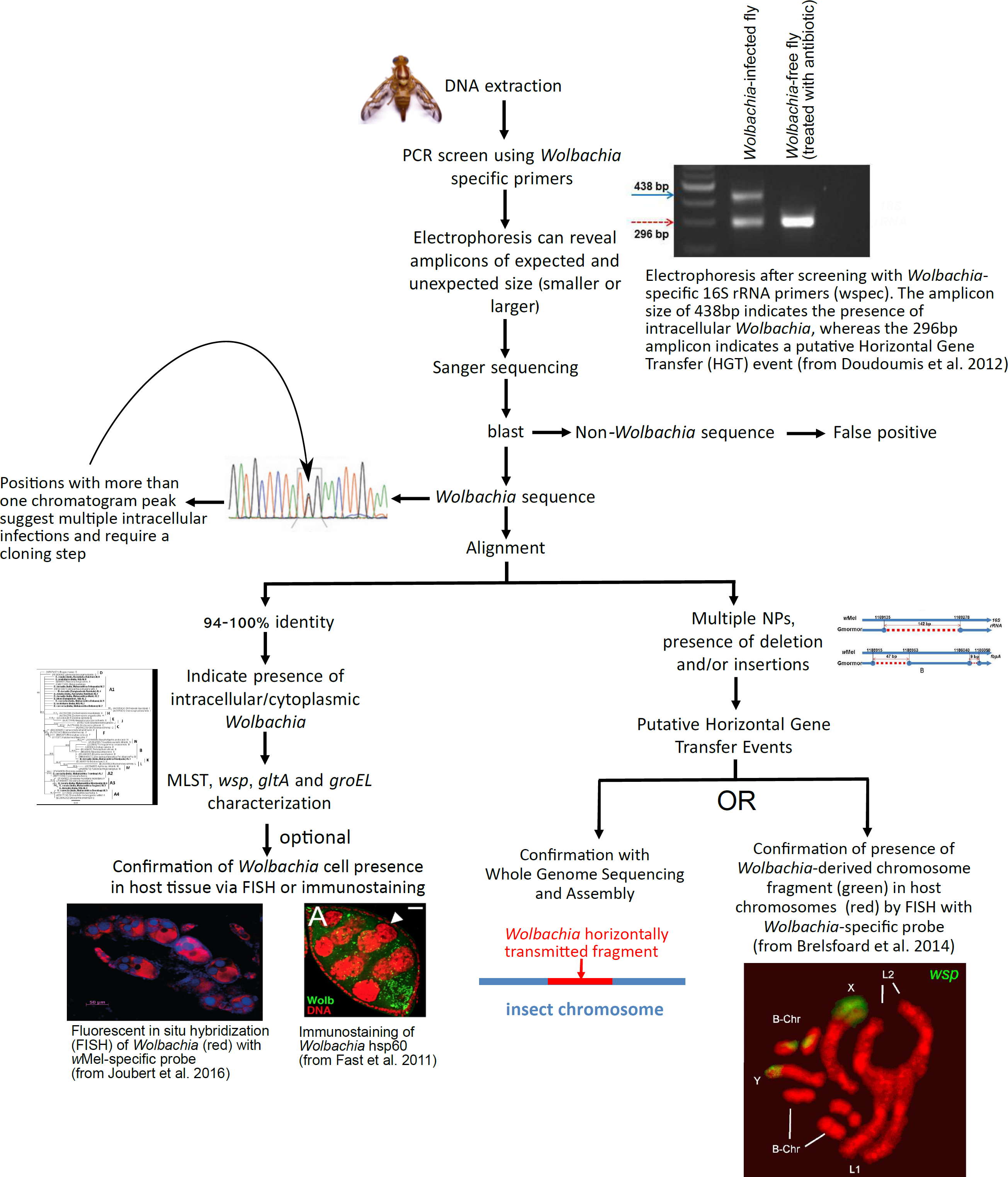
Recommended steps to screen for *Wolbachia* infections in tephritids and other arthropods. A PCR is performed with *Wolbachia*-specific primers on DNA isolated from whole, or parts of (e.g. abdomens), insect. Agarose gel electrophoresis of the PCR products is used to determine whether the amplicon is of the expected size. Amplicons of expected size are directly sequenced (e.g. Sanger method). High sequence identity to other *Wolbachia* suggests *Wolbachia* infection. Clean chromatograms are consistent with a single *Wolbachia* strain. Otherwise, a cloning step to identify different *Wolbachia* alleles is required. Other genes are then amplified and sequenced for further genetic characterization of the strain. As an optional step, localization of *Wolbachia* cells within host tissues can be achieved by Fluorescent In Situ Hybridization (FISH) with *Wolbachia*-specific rRNA probe or immunolabeling with antibody specific to *Wolbachia* protein. An amplicon of an unexpected size might indicate the occurrence of a horizontally transmitted *Wolbachia* genome fragment to the insect chromosome, rather than a current infection. Similarly, multiple nucleotide polymorphisms (NP) or insertions/deletions, compared to known strains, are suggestive of *Wolbachia* pseudogenes (e.g. horizontally transferred to host genome). This can be further tested by *in situ* hybridization of *Wolbachia*-specific probe to host chromosomes, and/or by Whole Genome Sequencing of host [images from 98, 195, 251, 252]. Photo of fly (*Anastrepha obliqua*) by Fabiola Roque (ECOSUR-UT).

The two major shortcomings of utilizing solely PCR (or LAMP) to detect *Wolbachia* presence can be classified into the occurrence of false negatives and false positives. A false negative occurs when a specimen is infected by *Wolbachia*, yet the screening approach fails to detect its presence. The efficiency of the PCR can be affected by the presence of inhibitors [91, 92], by low concentration/poor quality of the target DNA molecule, as well as type and concentration of the polymerase and other PCR reagents. At the very least, negative *Wolbachia* detection PCRs should be validated by evaluating the quality of the DNA extract, through positive amplification of a host-encoded gene (e.g. the mitochondrial Cytochrome Oxidase subunit I or single-copy nuclear genes). Several higher sensitivity approaches have been devised, particularly for low-titer infections, such as: long PCR [93]; nested PCR [94]; quantitative PCR [95]; or the design of alternative and/or more specific primers, including the use of *Wolbachia* multi-copy genes as PCR targets [96]. These methods, however, have not been widely implemented, likely due to the higher effort or cost involved.

False positives occur when a specimen not harboring *Wolbachia* is identified as *Wolbachia*-infected. Several instances have been reported where insect chromosomes carry *Wolbachia*-derived fragments, presumably from a horizontal gene transfer event that occurred at some point in the host lineage as the result of an active infection that was subsequently lost. The size of the horizontally-transmitted fragment can range from ca. 500 bp to the equivalent of an entire *Wolbachia* chromosome [97]. In some cases, entire *Wolbachia* chromosomes have been transferred more than once onto the same host genome [98, 99]. The range of hosts carrying *Wolbachia*-derived genome fragments is broad and includes several dipterans (tephritids, *Glossina morsitans* Westwood; *Drosophila spp.,* mosquitoes), other insects, as well as nematodes [97, 98, 100–102]. It is therefore desirable to corroborate PCR-based inferences with approaches that detect *Wolbachia* cells in host tissues. Such microscopy approaches can be based on nucleic acid hybridization [e.g. 103] or antibody-based detection of *Wolbachia* proteins [e.g. ftsZ; 104, and wsp; 105]. A major drawback of these methods is that they require substantial investment in time and equipment compared to PCR-based approaches. False positives can also occur if the primers targeted at *Wolbachia* turn out to amplify a fragment of the genome of the host (not derived from *Wolbachia*) or of another symbiont of the host. Such false positives are relatively easy to rule out upon sequencing and analysis of the amplified product. Finally, as with any PCR work, false positives can result from contamination of the specimen, the DNA template, or the PCR reagents. Thus, it is important to implement adequate sterile practices and negative controls.

The above approaches require destruction of specimens for DNA isolation or for tissue fixation. As a rapid and non-destructive alternative, Near-Infrared Spectroscopy (NIR) has been developed for identification of specimens infected with *Wolbachia*, including the distinction of two different *Wolbachia* strains [106]. This method, however, requires standardization according to species, sex, age, or any other condition that may affect absorbance, and is not 100% efficient. To our knowledge, this method has not been employed to assess *Wolbachia* infection in tephritids.

### 2.2 Methods to taxonomically characterize Wolbachia strains

The main evolutionary lineages of *Wolbachia* are assigned to “supergroups” [107]. Sixteen supergroups have been recognized to date [108, 109]. Supergroups A and B are widespread in arthropods and are common reproductive manipulators [39, 110]. Supergroups C and D are obligate mutualists of filarial nematodes, whereas supergroup F is found in both arthropods and nematodes [111]. Other supergroups have more restricted host distributions [112]. *Wolbachia* are generally compared and classified on the basis of Multilocus Sequence Typing (MLST) systems [110, 113]. The most commonly used MLST is based on the PCR amplification of fragments of five ubiquitous genes: *cox*A, *fbp*A, *fts*Z, *hcp*A and *ga*tB. However, this MLST system has limitations, in that not all genes are readily amplified in all *Wolbachia* strains, and it fails to distinguish among very closely related strains [109, 112]. Several additional genes commonly amplified and reported are the 16S *rRNA*, *gro*EL, *glt*A, and the *wolbachia surface protein* (*wsp*) [107, 112, 114, 115]. The *wsp* gene is highly variable and shows evidence of intragenic recombination [116, 117]. An MLST database (https://pubmlst.org/wolbachia/) is available to compare sequences of alleles for the five MLST loci and the *wsp* gene. Upon submission to the MLST, new alleles for the *wsp* and for each of the MLST loci are assigned a unique number. A *Wolbachia* sequence type (ST) is defined on the basis of allele combinations, with each allele combination assigned a unique ST number.

Hosts can be infected by one or more distinct strains of *Wolbachia*. Traditionally, direct Sanger sequencing of PCR products that resulted in sequences with ambiguous base pairs would be subjected to cloning followed by sequencing. The allele intersection analysis method [AIA; 118] can then be used to assign MSLT alleles to *Wolbachia* strains, but it requires a priori knowledge on the number of strains present. AIA identifies pairs of multiply infected individuals that share *Wolbachia* and differ by only one strain. Alternative approaches to circumvent cloning include the use of strain-specific primers [e.g. for the wsp gene; 107], or of high throughput sequencing approaches (e.g., Illumina HiSeq or Illumina MiSeq) to sequence MLST or other marker PCR amplicons [e.g. 119, 120]. Primer bias, however, where the fragment of one *Wolbachia* strain is preferentially amplified over the other, has been reported [118], such that presence of certain *Wolbachia* strains might be missed.

Use of the MLST system alone has two major drawbacks. First, strains of *Wolbachia* sharing identical MLST or *wsp* alleles can differ from each other at other loci [113, 121]. Secondly, the MLST, 16S *rRNA*, and *wsp* loci contain limited phylogenetic signal for inferring relationships within *Wolbachia* supergroups [109]. Therefore, to assess such intra-ST variation and to infer evolutionary relationships among closely related *Wolbachia* strains, additional (more variable) loci must be evaluated. The multiple locus variable number tandem repeat analysis developed by Riegler et al. [121] allows distinction of closely related *Wolbachia* strains based on PCR and gel electrophoresis.

Whole genome sequencing represents a powerful approach to distinguish closely related *Wolbachia* strains, infer their evolutionary relationships, test for recombination, and identify genes of interest [e.g. 47, 122]. Due to its fastidious nature [but see 123 for a recent breakthrough] and occurrence of repetitive elements, genome sequencing and assembly of *Wolbachia* (and other host-associated uncultivable bacteria) has proven difficult. Recent advances, particularly those based on targeted hybrid enrichment [124] prior to high-throughput sequencing [125, 126] have been successfully applied to *Wolbachia* for short-read technologies [i.e., Illumina; 127, 128]. The combination of targeted hybrid capture and long-read technologies, such as Pacific Biosciences’ Single Molecule Real-Time [e.g. 129] or Oxford Nanopore Technologies’ platforms is expected to greatly advance *Wolbachia* genomics research.

### 2.3 Methods to functionally characterize Wolbachia strains

A major challenge to investigating the effects of *Wolbachia* on a host is to generate *Wolbachia*-present and *Wolbachia*-free treatments while controlling for host genetic background. The challenge stems from the difficulty of adding or removing *Wolbachia* to/from particular hosts. Addition of *Wolbachia* to a particular host background can be achieved by transinfection [reviewed in 130]. Because the vertical transmission of *Wolbachia* appears to be dependent on its close association to the host germline, successful artificial transfer of *Wolbachia* typically relies on injection of cytoplasm from a donor egg [but see 131] or early embryo into a recipient embryo via microinjection [reviewed in 130]. The success rate of the transinfection procedure is generally very low; in tephritids it is 0–0.09% (calculated as the proportion of injected embryos that emerged as *Wolbachia*-infected adult females that transmitted *Wolbachia* to offspring [132, 133, 134; Morán-Aceves et al., unpublished data]). The low success rate is generally a result of low survival of injected embryos, the low proportion of *Wolbachia*-positive survivors, some of which do not transmit *Wolbachia* to their offspring.

Intra-species (or between sibling species) transfer of *Wolbachia* to a particular host nuclear background can also be achieved through introgression, whereby males of the desired background are repeatedly backcrossed with *Wolbachia*-infected females [e.g. 135, 136]. Under this scheme, after eight generations of consistent backcrossing, ˜99.6% of the host nuclear background is expected to have been replaced (Fig. 1d). The drawback of this approach is that the mitochondrial genome will not be replaced. Therefore, the effects of mitochondrial type and *Wolbachia* infection cannot be separated.

Due to less than perfect transmission, passive loss of *Wolbachia* in certain host individuals may be used to obtain *Wolbachia*-free and *Wolbachia*-infected hosts of equivalent genetic background. *Wolbachia* removal has also been achieved by “extreme” temperature treatment [e.g. 30°C; 137]. The most common way of removing *Wolbachia*, however, is achieved via antibiotic treatment, but several potential biases must be addressed [reviewed by 138]. Antibiotic treatment is likely to alter the microbiota, other than *Wolbachia*, associated with the host. In addition, antibiotics may affect the host in a microbe-independent manner. For instance, antibiotic treatment can affect host mitochondria [139], which in turn can reduce host fitness. A common practice to circumvent these problems is to wait several generations after antibiotic treatment, and to promote “restoration” of the host’s pre-antibiotic microbiota, excluding *Wolbachia* (e.g. exposing the insects to the feces of non-treated individuals). *Wolbachia* does not appear to be efficiently transmitted via ingestion [e.g. 140], but see discussion on horizontal transmission routes below. It is important to monitor *Wolbachia* infection status of antibiotic-treated host strains, because antibiotics may not always fully remove infection. Instead, they may reduce *Wolbachia* densities to non-detectable levels in one or few generations [138]; this has been our experience in both *Anastrepha* and *Drosophila* (unpublished data).

Unidirectional CI is tested by comparing the embryo hatching rates of the CI cross (uninfected female X infected male) to that of one or more control crosses. For testing bidirectional CI, the reciprocal crosses of hosts infected by the different *Wolbachia* strains are assessed. A significantly lower embryo hatching rate of the CI cross(es) compared to that of the control cross(es) constitutes evidence of CI. CI can be partial or complete (100% embryo failure). As with any fitness assay, care must be taken to prevent potential biases, including crowding and age effects; male age can influence CI [33, 81, 141]. Adequate assessment of fertilization must be performed to ensure that failed embryos are not confused with unfertilized eggs. This may require testing for insemination of females that produce no larval progeny [e.g. 134, 142], or exclusion of females that predominantly lay unfertilized eggs, such as old *D. melanogaster* virgin females [143].

## 3 Wolbachia in tephritids

### 3.1 Taxonomic distribution of Wolbachia-tephritid associations

Based mostly on PCR and sequencing approaches, ˜66% of ˜86 tephritid species screened have at least one record of positive *Wolbachia* infection (excluding pseudogenes) in laboratory and natural populations (Table S1; only supergroups A and B have been found in tephritids). For the genus *Anastrepha*, all but one species (*A. ludens*) of 17 screened to date harbor *Wolbachia* [142, 144–148, 149; and this study, 150]. Most *Anastrepha* species harbor *Wolbachia* strains assigned to supergroup A. *Anastrepha striata* Schiner and *Anastrepha serpentina* Wiedemann, however, harbor supergroup B in southern Mexico [147; and this study] and supergroup A in Brazil [146]. Up to three *Wolbachia* sequence types have been detected per locality within morphotypes of the *A. fraterculus* complex [142, 150], but co-infection of a single individual is generally not observed [except for one report in *A. fraterculus*; 151].

Of the ˜49 species of *Bactrocera* that have been examined, ˜14 are reported to harbor *Wolbachia* (supergroup A and/or B) and three (*Bactrocera peninsularis* Drew & Hancock, *Bactrocera perkinsi* Drew & Hancock and *Bactrocera nigrofemoralis* White & Tsuruta) carry what appear to be *Wolbachia*-derived pseudogenes, but not active infections [101, 132, 152–154]. There is also the case of *Bactrocera zonata* Saunders, *B. dorsalis*, and *Bactrocera correcta* Bezzi that have been found to carry both active infections (cytoplasmic) and pseudogenized *Wolbachia* sequences (Asimakis et al., in review). Up to five *Wolbachia* strains have been reported in a single individual of *Bactrocera ascita* Hardy [152], and double/multi infections have been reported in individuals of the following five *Bactrocera* species in Australia: *Bactrocera bryoniae* Tryon; *Bactrocera decurtans* May; *Bactrocera frauenfeldi* Schiner; *Bactrocera neohumeralis* Hardy; and *Bactrocera strigifinis* Walker [101, 153]. Within the genus *Bactrocera*, polyphagous species are more likely to harbor *Wolbachia* compared to stenophagous or monophagous ones [154].

For the genus *Ceratitis*, two species have been screened for *Wolbachia.* No evidence of *Wolbachia* was found in *Ceratitis fasciventris* Bezzi. Also, no evidence of infection was found in several populations of *C. capitata*, the Mediterranean fruit fly (medfly), in the early ‘90s [155]. PCR amplification and sequencing of the 16S *rRNA* gene in several field and lab specimens of *C. capitata* from Brazil suggested infection with *Wolbachia* supergroup A [156]. However, recent thorough surveys of wild populations and lab colonies indicate that *Wolbachia* is absent in *C. capitata* from numerous localities in different continents (Table S1). Two out of the six species of *Dacus* examined to date are reported to harbor *Wolbachia*: *Dacus axanus* Hering [153]; and *Dacus destillatoria* Bezzi [152]. *Wolbachia* has not been detected in the monotypic genus *Dirioxa* [101].

*Wolbachia* is reported in the four species of *Rhagoletis* examined to date: *Rhagoletis cerasi* L.; *Rhagoletis pomonella* Walsh; *Rhagoletis cingulata* Loew; and *Rhagoletis completa* Cresson [133, 157–164]. Both A and B supergroups are found in *R. cerasi* and *R. completa*, including a putative A-B recombinant strain, and co-infections are common (e.g. *R. cerasi* and *R. pomonella*). In *Zeugodacus* (formerly *Bactrocera*), both *Z. cucurbitae* and *Z. diversa* are reported to harbor *Wolbachia* [152, 154; Asimakis et al. in review].

### 3.2 Wolbachia prevalence in tephritids (in time/space)

Numerous studies report *Wolbachia* infection frequencies (or data from which this measure can be estimated) in natural populations of tephritids. Few of these studies, however, have adequate sample sizes for such inferences (e.g. many such studies are based on 10 or fewer individuals). Notwithstanding, inferred *Wolbachia* prevalence in tephritid populations is highly variable. In *Anastrepha*, ˜10 species harbor at least one population with prevalence ˜100%, whereas populations of three species reported lower frequencies (e.g. 88%, 51–60%, and 8.7%) (Table S1). In *Bactrocera*, one population of *B. caudata* had a 100% prevalence, whereas all other species with positive *Wolbachia* results exhibited low prevalence.

The best studied tephritid system in terms of spatial and temporal variation in *Wolbachia* prevalence is probably that of *R. cerasi* in Europe, which was surveyed over a ˜15 year period in 59 localities [163]. In an early (1998) survey of Riegler and Stauffer [165], all European *R. cerasi* individuals were infected by one strain (*w*Cer1), most central and southern European populations harbored an additional strain *w*Cer2 (i.e., were co-infected), and at least an Italian population harbored *w*Cer4 [133]. A rapid spread of *w*Cer2 (a strain associated with cytoplasmic incompatibility) has been detected. Multiple infections in various combinations considering all five known *Wolbachia* strains have been revealed recently. Samples from Poland, Italy, and Austria, are infected with all five *w*Cer strains, those from Czech Republic and Portugal lack *w*Cer2 only, while the Swiss samples lack *w*Cer3 [162]. Recent analysis of 15 Greek, two German and one Russian population demonstrate fixation for *w*Cer1 in all *R. cerasi* populations and the presence of complex patterns of infections with the four of the five known *w*Cer strains (1, 2, 4, and 5) and the possible existence of new *Wolbachia* strains for the southernmost European *R. cerasi* population [i.e., Crete; 158]. Similarly, strain *w*Cin2 (which is identical to *w*Cer2 based on loci examined to date) is fixed in all populations of *R. cingulata*; a species native to the USA, but introduced into Europe at the end of the 20^th^century. Invasive populations in Europe harbor *w*Cin1 (identical to *w*Cer1 based on loci examined to date) at frequencies that vary over space and time (up to 61.5%), as a result of horizontal transfer (multiple events) from *R. cerasi* [163].

### 3.3 Phenotypic effects of Wolbachia in tephritids

Despite the numerous reports of *Wolbachia* in tephritids, the fitness consequences of such associations remain mostly unknown. The studies reporting phenotypic effects of *Wolbachia* have relied on transinfection and on antibiotic-curing; only two species of tephritids have been successfully transinfected with *Wolbachia* (Table 1). Evidence of *Wolbachia*-induced CI has been detected in three species of tephritids. Early studies [166, 167] identified reproductive incompatibilities in *R. cerasi* that were later attributed to the *Wolbachia* strain *w*Cer2 [100% embryonic mortality in the CI cross; 165]. Artificially transferred *Wolbachia* (strains *w*Cer2 and *w*Cer4) originally from *R. cerasi* to *C. capitata* also resulted in strong CI (100% embryonic mortality). *w*Cer2 in two genetic backgrounds of *B. oleae* resulted in strong CI as well [132]. In addition, *w*Cer2 and *w*Cer4 are bi-directionally incompatible in *C. capitata* [133, 134].

**Table 1.** Successful and unsuccessful *Wolbachia* transfection attempts in tephritids.

| **ID of successfully transfected tephritid strain** | **Donor species /strain** | **Recipient species (and strain)** | ***Wolbachia* strain** | **Reference** |
| --- | --- | --- | --- | --- |
| *C. capitata* WolMed 88.6 | *R. cerasi* | *C. capitata* Benakeio strain | wCer2 | [133] |
| *C. capitata* WolMed S10.3 | *R. cerasi* | *C. capitata* Benakeio strain | wCer4 | [133] |
| *C. capitata* VIENNA 8-E88 | *C. capitata* WolMed 88.6 | *C. capitata* VIENNA 8 Genetic Sexing Strain (GSS) | wCer2 | [134] |
|  | *C. capitata* VIENNA 8-E88 | *Bactrocera oleae* | wCer2 | [132] |
| N/A (unsuccessful) | *A. striata* | *A. ludens* | wAstriB | (unpublished) |

In addition to CI, *Wolbachia*-infected *C. capitata* females (Benakeio strain) exhibit higher embryonic mortality (17–32% in crosses with *Wolbachia*-free males and 65–67% in crosses with *Wolbachia*-infected males) than their *Wolbachia*-free counterparts crossed with *Wolbachia*-free males (12% embryonic mortality). Therefore, it appears that *w*Cer2 and *w*Cer4 have additional fertility effects on medfly females, other than CI. It is also possible that these *Wolbachia* strains can only partially rescue the modification that they induce in sperm [133]. A similar pattern is reported in the Vienna 8 genetic sexing strain (GSS) infected with *w*Cer2 [134]. The *w*Cer2 also causes increased embryo death in non-CI crosses of *B. oleae* [132].

Two studies conducted several years apart [168–170] examined the effects of a single *Wolbachia* strain (*w*Cer2) on fitness components of two *C. capitata* genotypes (i.e., Benakeio and Vienna 8 GSS laboratory lines), as well as the effects of two different *Wolbachia* strains (*w*Cer2 and *w*Cer4) on a single medfly genotype (Benakeio). The following general patterns emerged (exceptions noted): (a) *Wolbachia* causes higher egg-to-larva mortality; (b) *Wolbachia* causes higher egg-to-adult mortality [exception: Vienna 8 GSS + wCer2 in 170]; (c) *Wolbachia* shortens egg-to-adult development time [exception: Benakeio + wCer2 in 168, 169]. In addition, Sarakatsanou et al. [170] found that *Wolbachia* shortens both male and female adult lifespan (exception: males of Vienna 8 GSS + *w*Cer2), and reduces life time female net fecundity. However, Kyritsis G.A [168], Kyritsis G.A. et al. [169] reported no effects of *Wolbachia* infection on adult lifespan, and a reduced fecundity in the case of *w*Cer4 infection only. Even though *w*Cer2 and *w*Cer4 in general tended to have consistent effects on medfly, the magnitude of their effects differed. Collectively, the results from these studies indicate that the effect of *Wolbachia* infection on life history traits depends both on the *C. capitata* genetic background and on the *Wolbachia* strain. Furthermore, inconsistencies between the two studies might be indicative of evolution of the host and/or *Wolbachia* strain during that period. Adult flight ability and longevity under stress conditions also appear to be determined by the interaction of *Wolbachia* strain and medfly genotype [168, 169].

A recent study by Conte et al. [142] examined the phenotypic effects induced by two *Wolbachia* strains native to *A. fraterculus* (sp1). No evidence of bidirectional cytoplasmic incompatibility was detected in reciprocal crosses. Ribeiro [137] reported evidence consistent with CI caused by *Wolbachia* in *A. obliqua* and in *“A. fraterculus sp. 1”*, which according to *wsp* sequences, are identical. Nonetheless, confounding effects of the treatment to remove *Wolbachia* (removed by exposure of pupae to 30 °C) or other potential biases cannot be ruled out, as all intraspecific crosses involving at least one cured parent resulted in much lower (< 30%) embryo hatching than the intraspecific crosses involving both infected parents (66 and 81% embryo hatching). Whether *Wolbachia* influences any other fitness aspects of tephritids, including its interactions with other microbes, has not been evaluated.

Very little is known regarding the effect of *Wolbachia* on tephritid behavior. Recent tests demonstrated that *Wolbachia* infection affected male sexual competitiveness. Different *Wolbachia* strains (*w*Cer2 and *w*Cer4) exerted differential impact on males mating competitiveness, and a single strain (*w*Cer2) had different impact on different medfly genotypes (Benakeio and Vienna 8 GSS laboratory lines) [168, 169].

### 3.4 Modes of horizontal transmission of Wolbachia between tephritid hosts

Considering the dynamics of *Wolbachia* associated with arthropods in general, at the population level, *Wolbachia* appears to be predominantly maintained by vertical transmission. Above the species level, however, the lack of congruence between the host and symbiont phylogenetic trees implies that *Wolbachia* horizontal transfers and extinctions are common and underlie its widespread taxonomic and geographic distribution [171].

The possible routes by which *Wolbachia* may be horizontally acquired by a new host can generally be classified as via ingestion or via a vector. In both cases, to become established as a stable cytoplasmically inherited infection, *Wolbachia* must cross one or more cell types or tissues. For example, if *Wolbachia* invaded the host hemolymph directly as a result of a vector (e.g. parasitoid wasp or ectoparasitic mite), it would have to invade the egg during oogenesis. Similarly, if *Wolbachia* were acquired via ingestion (e.g. as a result of scavenging), it would have to cross the gut into the hemolymph, before it invaded the egg. Support for the above routes comes from studies reporting: (a) that *Wolbachia* can remain viable in an extracellular environment and infect mosquito cell lines, as well as ovaries and testes that are maintained *ex vivo* [172, 173]; (b) that *Wolbachia* cells injected into *Drosophila* hemolymph reach the germline after crossing multiple somatic tissues [131]; (c) that *Wolbachia* can move between parasitic wasp larvae (*Trichogramma*) sharing the same host egg, and achieve vertical transmission [174]; and (d) that parasitic wasps of the white fly, *Bemicia tabaci* (Gennadius) can transfer *Wolbachia* from an infected to a naïve host, as a result of non-lethal probing (i.e., probing without oviposition), whereby the parasitoid ovipositor or mouthparts function as a “dirty needle” [175].

No direct evidence of *Wolbachia* transmission via parasitoids exists in tephritids, but sharing of *Wolbachia* strains between a parasitoid and several sympatric tephritids [144, 153] is consistent with parasitoid-mediated transmission, or transmission from tephritid host to parasitoid [176]. The potential for horizontal transfer of *Wolbachia* among tephritids via parasitoids is high, due to the multiple instances where a single parasitoid utilizes several different tephritid host species [177–179].

*Wolbachia* may invade a new host species via introgressive hybridization between two host species. This mechanism would also transfer mitochondria from the infected to the uninfected species nuclear background, akin to the artificial backcrossing approach described above (Fig. 1d). Ability of tephritids to hybridize in the lab has been reported in numerous species (Table 2), and hybridization in nature has been documented in *B. dorsalis/B. carambolae* [180], members of the *C.* FAR complex [181], *R. cingulata/R. cerasi* in Europe [182]. Thus, there is potential for wild tephritid populations to acquire *Wolbachia* infections via hybridization.

**Table 2.** Representative tephritid genera where hybridization between one or more species has been reported.

| **Tephritid genera containing species that can hybridize** | **Reference(s)** |
| --- | --- |
| *Bactrocera* | [231-236] |
| *Ceratitis* | [237] |
| *Anastrepha* | [238-244] |
| *Rhagoletis* | [245-249] |
| *Eurosta* | [250] |

## 4 Considerations for Wolbachia-based IIT in tephritids

### 4.1 The need for genetic sexing strains (GSS)

In both SIT and IIT, the purpose of releasing insects is to reduce the fertility of the wild population, as a result of mating by wild females with sterile or sterility-inducing males, respectively. SIT is most effective when only males are produced and released [183, 184], but release of a small percentage of females, which are typically sterilized with lower doses than males, is not considered overly detrimental to the program. The release of only males in a large-scale operation can be accomplished by either killing female zygotes during development or by selectively removing them from the mass-reared population prior to release [185, 186]. In most tephritids, male sex is determined by the presence of the maleness factor on the Y chromosome [187]. GSS based on male-linked (e.g. Y chromosome – autosome (Y;A)) translocations have been developed in a few species to produce conditional female lethality (e.g. temperature sensitive lethality during embryonic development) or a visual sex marker (e.g. pupal color). Unfortunately, despite substantial effort, GSS are still lacking for most tephritid pests. The recent development of CRISPR/Cas9-mediated mutagenesis in tephritids, however, might enable a faster development of tephritid GSS [188, see reviews by 189, 190].

Whereas the use of GSS is **beneficial** to SIT programs, it is an **absolute necessity** for IIT programs, unless mitigating mechanisms can be implemented. In an IIT operation, the release of even a few *Wolbachia*-infected females could be disastrous, because these females would be fertile with both wild and released males (which would carry a compatible *Wolbachia* strain). It is unlikely that any sexing system will achieve zero release of females. Therefore, a mechanism that guarantees that any released females are sterile is necessary. If female sterility is induced with substantially lower radiation doses than male sterility, then radiation could be used to render females sterile without the detrimental effects of the higher doses required to sterilize males, and this is one of the advantages of combining the SIT with the IIT approach [17, 21, 22, 191–193].

### 4.2 Choice and evaluation of Wolbachia strains

The goal is to generate a mass-reared colony that is infected with a mod^+^/resc^+^ *Wolbachia* strain that causes strong incompatibility with the target pest population. The target pest population will be *Wolbachia*-free (e.g. most *C. capitata* wild populations and all *A. ludens* populations) or harbor one or more *Wolbachia* strains. Below we list recommendations regarding the selection and evaluation of the *Wolbachia* strain to use in IIT and in SIT/IIT approaches. The target population and the donor colony should be thoroughly screened for *Wolbachia*, ideally with the higher sensitivity methods described in Section 2.1, to detect low-titer and multi-strain infections. The *Wolbachia* strain selected should be incompatible with those found in the target population. The selected *Wolbachia* strain should be artificially transferred to one or more lab colonies, representative of the genetic background of the target pest population. Most cases of successful establishment of stable transinfected insect lines have relied on embryonic microinjection [130]. Introgressive backcrossing might be feasible in scenarios where geographically isolated populations of the same target species harbor distinct *Wolbachia* strains (e.g. *A. striata* in Mexico vs. *A. striata* in Brazil; Table S1). A thorough biological characterization of the artificial host-*Wolbachia* association should be conducted, as both host background and *Wolbachia* strain are important determinants of CI expression and other relevant fitness parameters [17, see also 169, reviewed in 194]. The main desired characteristics of the association are: strong induction of CI; no rescue by *Wolbachia* strain(s) present in target population; few or no fitness costs for parameters relevant to the program. These fitness parameters can be classified into those related to a cost-effective mass production (e.g. female fecundity including embryo hatching success) and those related to the success of released males (e.g. mating and sperm competitive ability). Some host-*Wolbachia* combinations result in higher female fecundity, such as *Drosophila simulans* after many generations [60] and *Drosophila mauritiana* [195]. In contrast, other host-*Wolbachia* combinations result in lower fertility [e.g. low embryo success in *C. capitata* and *B. oleae*; 132, 133, 134]. *Wolbachia* could affect male mating success by influencing assortative mating; a phenomenon detected in some studies of *Drosophila* [e.g. 196, 197], but not others [e.g. 198, 199, 200]. Such influence of *Wolbachia* on mating preferences was questioned [201] on the basis of evidence that gut microbiota influence assortative mating in *Drosophila* [198, 201, 202], a finding that itself has been questioned recently [203]. In addition, at least one case has been reported where sperm from *Wolbachia*-infected males were less competitive [79]. Similarly, *Wolbachia*-infected *D. simulans* produce fewer sperm [83]. All of the above parameters should be evaluated under relevant conditions known to interact with *Wolbachia*, such as temperature and nutrition [reviewed in 17, 204, e.g. 205], interaction with other microorganisms [e.g. 206, 207], as well as male age and mating status [e.g. 208, 209].

### 4.3 Other considerations

#### 4.3.1 Species recalcitrant to Wolbachia?

Certain species or clades appear to be “resistant” to *Wolbachia* infection, based on their lack of infection in nature and the failure to achieve stable transfections. The reasons are unknown, but could involve host and/or bacterial factors. For example, none of the members of the diverse *repleta* species group of *Drosophila*, comprised mostly of cactophilic flies [210], has ever been found to harbor *Wolbachia* [211]. Similarly, due to numerous failed transinfection attempts, and the lack of natural infection in wild *Anopheles* mosquitos, this genus was regarded impervious to *Wolbachia* [reviewed in 130]. This view has been challenged by the recent successful establishment *Wolbachia-*transfected *Anopheles stephensi* Liston [62], and the recent discovery of a natural stable *Wolbachia* infection in *Anopheles coluzzii* Coetzee & Wilkerson [212]. The lack of natural infections and transinfection failure in *A. ludens* may reflect a general refractoriness to *Wolbachia*. Nonetheless, initial attempts to transinfect *C. capitata* also failed and transfection with *Wolbachia* was attained later on with different *Wolbachia* strains [133]. Hence, transinfection attempts with additional *Wolbachia* strains may result in successful and stable infection in *A. ludens* as well.

#### 4.3.2 Potential for target populations to become resistant to sterile males

There are two ways in which a target population may become resistant to the effects of release of *Wolbachia*-infected males. The first is endosymbiont-based, whereby the target population may acquire (e.g. via horizontal transmission) a *Wolbachia* strain that can rescue the modification (sterility) induced by the strain present in the released males. Generally, such acquisition of a *Wolbachia* strain during the relatively short lifespan of a release program is highly unlikely. Nonetheless, knowledge on the *Wolbachia* infection status and strain identity of interacting species, such as other fruit flies sharing the same host plant and parasitoids, might aid in the selection of *Wolbachia* strains that are unlikely to be compatible with strains that can potentially be horizontally acquired by the target population. Permanent screening of wild flies from the target population could provide valuable information in order to foresee potential lack of effectiveness of the method.

The second mechanism is host-based, whereby pre- or post-mating selection on wild females to avoid or reduce fertilization by incompatible sperm [reviewed by 213], acts on standing (or *de novo*) genetic variation. Evidence consistent with influence of *Wolbachia* on premating mechanisms comes from the observation that females and males of *Drosophila paulistorum* Dobzhansky and Pavan exhibit assortative mating according to the *Wolbachia* strain they harbor [197]. In addition, treatment with antibiotic (which removed *Wolbachia*) decreases mate discrimination in *D. melanogaster* [196]. The evolution of resistance to mating with mass-reared males by wild females can be potentially minimized by frequently refreshing the genetic background of the mass-reared strain [214–216], which is a routine process in mass-rearing programs [217]. Nonetheless, if the basis for mate discrimination were solely determined by *Wolbachia* infection state (e.g. if females could distinguish *Wolbachia*-infected vs. *Wolbachia-*uninfected males solely on the basis of a *Wolbachia*-encoded factor), refreshing the fly genetic background of mass-reared strain is unlikely to slow down the evolution of resistance to released males in the target population.

Several lines of evidence are consistent with the influence of *Wolbachia* infection on post-mating mechanisms. The existence of genetic incompatibility is predicted to favor polyandry (multiple mating by females) as a female strategy to minimize the probability of her eggs being fertilized by sperm from incompatible males [218]. Consistent with this prediction, uninfected *D. simulans* females remate sooner than *Wolbachia*-infected females [219]. Furthermore, *Wolbachia* modifies the length of the spermathecal duct of females of the cricket *Allonemobius socius* Scudder [220], which in turn may afford the female greater control on the outcome of sperm competition [e.g. D. melanogaster; 221]. Finally, the fact that host background can influence the CI phenotype [reviewed by 17], suggests that target populations may have genetic variants that are more resistant to CI, which could increase in frequency as a result of the strong selection exerted by the massive release of *Wolbachia*-infected males.

#### 4.3.3 Potential alternative ways of implementing Wolbachia-based approaches

The recent identification of *Wolbachia* “CI genes” offers potential alternative ways of harnessing reproductive incompatibility in control of pest tephritids. First, to identify strains with the desired characteristics, at least ability to induce CI, a productive endeavor might be to search for CI loci in the genomes of candidate strains being considered for IIT, prior to artificial transfer efforts. A candidate *Wolbachia* strain that lacks CI loci homologues, or that contains CI loci homologues that are highly similar to (and thus potentially compatible with) strains present in target population, should be avoided. Secondly, it may be possible in the future to genetically engineer *Wolbachia* strains with the desired characteristics (e.g. one or more specific CI operons) for IIT programs, or to replace strains used previously in a control program, as a means of addressing resistance phenomena [222]. Finally, a thorough understanding of the CI mechanism might enable the development of IIT based on *Wolbachia* transgenes, rather than *Wolbachia* infection. This might be particularly helpful in the control of species that are resistant to *Wolbachia* infection. Nonetheless, the release of such genetically modified insects might not be feasible due to regulatory hurdles and lack of public acceptance.

It has recently been shown that some *Wolbachia* strains can provide protection against major pathogens and parasites of insects, including RNA viruses and bacteria [66, 71, 223, 224]. It is very common for pathogens to appear in rearing facilities. Thus, if a *Wolbachia* strain could simultaneously cause strong CI and protect against a one or more pathogens (e.g. RNA virus), this would have multiple benefits in an operational *Wolbachia*-based population suppression program. Furthermore, a *Wolbachia* strain that does not induce (strong) CI, but protects against pathogens might be desirable in a program that does not rely on CI (e.g. SIT) for population suppression. *Wolbachia-*mediated pathogen protection would enable high production and quality levels, thereby contributing to a cost-effective and sustainable insect pest management program.

#### 4.3.4 Potential influence of other heritable symbionts

Multiple studies have revealed that although *Wolbachia* appears to be the dominant facultative heritable symbiont of arthropods, numerous other diverse bacteria (e.g. *Spiroplasma*, *Arsenophonus*, *Rickettsia,* and *Cardinium*) form such associations with insects, causing a diversity of reproductive and non-reproductive phenotypes [reviewed in 225, 226, 227]. Despite the long-standing recognition that “*Wolbachia* do not walk alone” [228], many studies of *Wolbachia* fail to rule out the association of their study organism with other facultative heritable symbionts. Even intensely studied groups in terms of heritable symbionts, such as tsetse flies (genus *Glossina*), can yield surprises of bacterial associates [e.g. the recent discovery of Spiroplasma in two species of Glossina; 229]. With few exceptions [142, 147; Asimakis et al., unpublished, 230], research on tephritid facultative heritable bacteria has not examined the possibility of players other than *Wolbachia*. Therefore, we urge that such research include screens for other symbionts, including viruses, protozoans, and fungi.

## 5 Conclusions

Given the widespread occurrence of *Wolbachia* in tephritids and its known fitness consequences in this group of dipterans and in other host taxa, *Wolbachia* is likely an influential component of tephritid ecology. Further exploration of *Wolbachia*-tephritid associations is expected to reveal a diversity of effects, as seen in more extensively study systems such as *Drosophila* and mosquitos. The recent exciting progress in understanding the basis of CI, and many other aspects of *Wolbachia* biology, should accelerate progress in the development of *Wolbachia*-based IIT for tephritid species. Nonetheless, IIT applications strongly depend on highly efficient and robust sexing strategies that are not yet available. Thus, IIT will most likely be used in combination with SIT since the use of radiation provides a failsafe mechanism for population suppression programs. In addition, the combined SIT/IIT approach may result in the use of reduced irradiation doses as well as enhanced pathogen protection and male mating competitiveness of mass-reared sterile insects. It is important, however, to not overlook the potential influence of other microbial players on the interaction between *Wolbachia* and its tephritid hosts.

## Acknowledgements

María de los Ángeles Palomeque-Rodas, Luz Verónica García-Fajardo, and Uriel Gallardo-Ortiz (ECOSUR-Unidad Tapachula) provided technical support. Carlos Cáceres (Insect Pest Control Laboratory, Joint FAO/IAEA Division of Nuclear Techniques in Food and Agriculture, Vienna, Austria) and Emilio Hernández-Ortiz (Programa Moscafrut, SAGARPA, SENASICA, IICA) provided materials. Funding was provided by the Joint FAO/IAEA Coordinated Research Project “Use of Symbiotic Bacteria to Reduce Mass-Rearing Costs and Increase Mating Success in Selected Fruit Pests in Support of SIT Application”. We thank two anonymous reviewers for their valuable suggestions.

---

**Supporting Table 1** (Excel spreadsheet)

Compilation of published and unpublished reports of screenings of *Wolbachia* (and other heritable bacteria) in pest Tephritidae. In our counts of species, *A. fraterculus* morphotypes [253] are regarded as separate species. Additional references not cited in main text but cited in this table [254–256].

